# Symmetric Inheritance of the Cytoplasmic Bacterium *Orientia tsutsugamushi* During Mitosis

**DOI:** 10.64898/2026.10.19.760126

**Authors:** Leana Keel, Buzz Baum, Jeanne Salje

## Abstract

For long-term stable residence within host cells, endosymbionts must evolve mechanisms to ensure their segregation into both daughter cells during cell division, and to coordinate their division with the host cell cycle. This regulation was also a prerequisite for the evolution of mitochondria, enabling their transition from free-living bacteria to an organelle. Here, we examine *Orientia tsutsugamushi* and *Rickettsia rhipicephali*, two obligate intracellular *Rickettsiales* that replicate directly in the host cytoplasm, during host mitosis. Using cell cycle synchronisation and immunofluorescence microscopy, we demonstrate that *O. tsutsugamushi* frequently localise near the spindle poles during mitosis, and are distributed between daughter cells in a manner dependent on its surface protein ScaC, that drives dynein-dependent motility of the bacterium along microtubules. In contrast, *R. rhipicephali*, which does not interact with microtubules, largely inhibits host mitosis and is randomly distributed in the few cells that divide. Overall, our findings reveal distinct strategies for mitotic adaptation among these two Rickettsiales and suggest that associations with the host’s cytoskeletal elements may be critical for near-equal distribution into daughter cells.

**Importance:** For long term endosymbiosis, intracellular bacteria must persist across host cell generations, but it remains unclear how pathogens like *Orientia tsutsugamushi,* adapt to host cell division to ensure their stable inheritance. Here, we report that *Orientia tsutsugamushi* localises to the mitotic spindle poles during host cell division and is distributed between daughter cells in a manner dependent on dynein-mediated transport along microtubules. These findings highlight how interactions between intracellular bacteria and the host cytoskeleton may reflect principles that enabled the evolution of stable endosymbiotic organelles such as mitochondria.

## Introduction

Endosymbiosis refers to a relationship where one organism lives inside another, typically establishing a long-term association. This interaction encompasses a spectrum from partially obligate to fully obligate, with varying degrees of inter-dependency. At the most extreme, both partners become mutually dependent and physically integrated, with the symbiont passed exclusively from parent to offspring. The most well-known examples of endosymbiosis are found in the origin of eukaryotic cells, as evidenced by the presence of mitochondria and chloroplasts^1^.

The establishment of a stable, long-term endosymbiosis within a eukaryotic cell requires the successful navigation of multiple challenges. Beyond the initial hurdle of host cell entry, a prospective endosymbiont must overcome host immune responses^2^ and align its metabolism with the environment inside of the host^3^. Another requirement is successful transmission, and an endosymbiont must develop mechanisms to ensure its passage to subsequent generations. At the cellular level this means that they must make sure that they are equally partitioned between both daughter cells to maintain the same bacterial load per host cell. Failure to do so would result in high variability in symbiont loads, where cells receiving too many symbionts could become overburdened, while those receiving too few or none risk losing the endosymbiont over time. Early proto-mitochondria faced similar pressures. To persist stably, they structurally integrated with the host cytoskeleton and tightly coordinated their replication with host mitotic processes^2^.

Bacteria that can only replicate within eukaryotic host cells are particularly relevant for understanding the establishment and maintenance of stable endosymbiosis. Such bacteria can be found within the alphaproteobacterial order Rickettsiales, including *Anaplasma*, *Ehrlichia*, *Rickettsia*, *Orientia*, and *Wolbachia*^4^. Their intracellular lifestyles vary considerably: members of the Anaplasmataceae, such as *Anaplasma, Ehrlichia* and *Wolbachia*, replicate within membrane-bound vacuoles^5–7^ whilst of the Rickettsiaceae family, such as *Rickettsia* and *Orientia*, escape the endolysosomal pathway shortly after entry and replicate directly in the host cytoplasm^4,8^. Within host cells, *Orientia tsutsugamushi* utilises the microtubule cytoskeleton and the motor protein dynein to traffic toward the microtubule organising center (MTOC), to the perinuclear region where bacterial division takes place^9^. This movement is mediated by the bacterial autotransporter protein ScaC, which recruits the dynein adaptors BICD1 and BICD2^10^. ScaC is sufficient to activate dynein-based motility, inducing conformational changes in BICD2 that promote the assembly of dynein-dynactin transport complexes^10^. In contrast, Spotted Fever Group (SFG) *Rickettsia* species rely on actin-based motility, rapidly inducing the polymerisation of host F-actin at one bacterial pole to generate comet-like tails^11,12^. These structures facilitate movement within the cytosol and support direct cell-to-cell spread. Because of their direct access to the host cytoplasm and association with cytoskeletal elements, *Rickettsia* and *Orientia* are valuable model systems for studying endosymbiosis.

Rickettsiales have evolved diverse strategies to adapt to the host cell cycle. *Anaplasma phagocytophilum* and *Ehrlichia chaffeensis* can manipulate or arrest the host cell cycle, stabilising the intracellular environment to support bacterial replication^13,14^. *Wolbachia pipientis* associate with centrosomes and exploit microtubule trafficking, ensuring even partitioning into daughter cells during embryogenesis^15,16^. Little is known about how *Orientia* and *Rickettsia* interact with the host cell cycle, although it has recently been shown that *O. tsutsugamushi* strain Ikeda induces cell cycle arrest in infected mammalian cells^17^.

To better understand the mechanisms that enable stable long-term endosymbiosis across multiple host generations, we investigated how cytoplasmic endosymbionts adapt to the challenges posed by host cell division. We first focused on *Orientia tsutsugamushi*, examining its behaviour during host mitosis in RPE1 human epithelial cells. Using immunofluorescence microscopy of synchronised cells, we sought to determine where these bacteria localise during mitosis and how they are partitioned between daughter cells, by disrupting their interactions with the host microtubule network. We then extended this analysis to *Rickettsia rhipicephali*, a cytoplasmic bacterium not known to interact with the host microtubule network, to compare how differences in host–cytoskeleton interactions influence bacterial localisation and segregation during host cell division.

## Results & discussion

*O. tsutsugamushi* localises to the mitotic spindle poles during host cell division and is distributed between daughter through dynein-mediated microtubule transport.

We first assessed whether *O. tsutsugamushi*–infected cells retain the ability to divide by quantifying the proportion of dividing mammalian cells in infected versus uninfected conditions following synchronisation using S-trityl-L-cysteine (STLC), an Eg5 inhibitor that blocks cell cycle progression at prometaphase Although *O. tsutsugamushi* infection modestly reduced the proportion of dividing cells (55.2% vs. 60.4% in controls; p = 0.013; Fig. 1A & 1B), infected cells remained able to enter mitosis.

**Figure 1:**
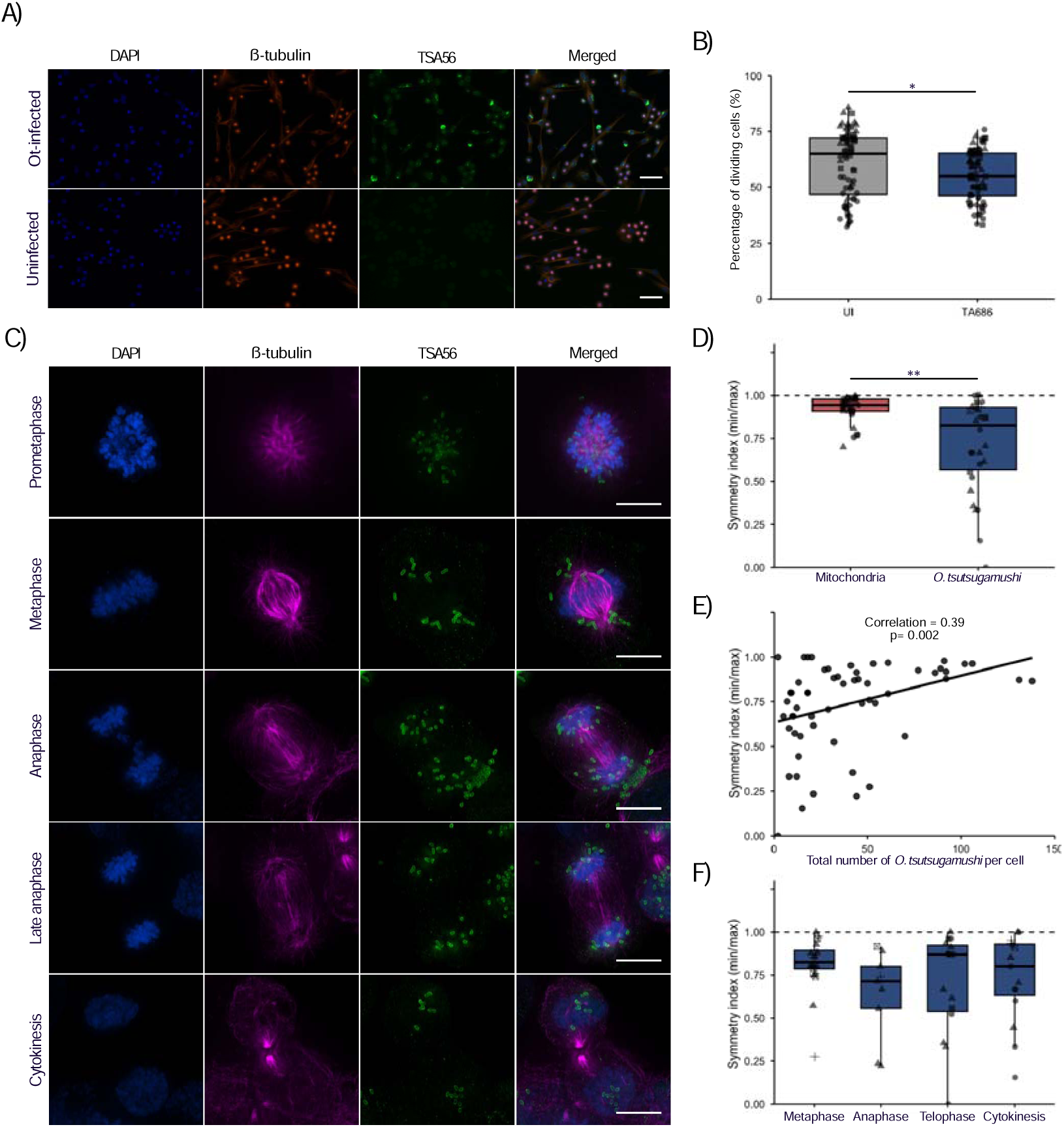
*O. tsutsugamushi* localises near spindle poles in dividing host cells, and exhibits a near-symmetric distribution of bacterial load between daughter cell. A. Representative image of Rpe1 cells infected for 6 days with Ot_TA686. Blue = DAPI (DNA), red = anti-β-tubulin antibody, green = anti-TSA56. B. Representative image of Rpe1 cells infected for 4 days with Rr. Blue = DAPI (DNA), red = anti-β-tubulin antibody, green = anti-OmpB. Images were acquired using a widefield microscope at 20X magnification. Scale bar = 50 μm. C. Quantification of data described in A. showing percentage of dividing cells in uninfected and infected samples. Each data point represents a single imaging field (n = 24 fields per replicate), acquired from three biological replicates. Given that an average of 95.88% ± 1.31% (SEM) of *Orientia*-exposed cells were infected across the repeats, any observed changes in division frequency can be confidently attributed to the presence of the bacteria. C. Representative images of Rpe1 cells infected for 6 days with *O. tsutsugamushi* strain Ot_TA686. Blue = DAPI (DNA), pink = anti-β-tubulin antibody, green = anti-TSA56. Images were acquired using a widefield microscope at 100X magnification. Scale bars represent 10 μm. D. Quantification of symmetry index (min/max bacterial number or mitochondrial signal in each daughter cell) of dividing cells in telophase or cytokinesis. A symmetry index of 1 indicates perfect symmetry. (Mitochondria: n=12, 10, 9 from 3 replicates; Ot_TA686: n=12, 11, 3, 2, 2 from 5 replicates). E. Scatter plots showing the relationship between total bacterial load and symmetry index in dividing cells from metaphase and later mitotic stages (Ot_TA686: n=12, 30, 3, 8, 6 from 5 replicates). Each point represents a single cell. The black line indicates the fitted linear regression. F. Quantification of symmetry index dividing cells in metaphase, telophase and cytokinesis (Metaphase: n = 12, 5, 3 from 3 replicates; Anaphase: n= 7, 1, 1; Telophase: n = 4, 7, 3, 1 from 4 replicates; Cytokinesis: n = 8, 4, 2 from 3 replicates). Boxplots display the median and interquartile range for each group. Statistical analysis was performed using two-sample Mann–Whitney tests for A. (Ot-infected vs. Uninfected), beta regression for D. (*O. tsutsugamushi* vs. mitochondria) and F (metaphase vs. anaphase, telophase and cytokinesis), with total bacterial number per cell included as a covariate, and Spearman’s rank correlation for E. Significance: **p < 0.01, *p < 0.05.

To visualise the intracellular distribution of *O. tsutsugamushi* during host mitosis, infected RPE1 cells were synchronised in prometaphase using STLC for 16-19 hours and were fixed 1 hour and 3 hours following drug washout to capture early and late mitotic stages. Antibodies against the *O. tsutsugamushi* outer membrane protein TSA56 were used to label the bacteria, and co-labelled with β-tubulin to accurately identify mitotic stages based on spindle morphology. During prometaphase, *O. tsutsugamushi* often localise in close proximity to the condensed nuclei (Fig. 1C). From metaphase onward, once the mitotic spindle is formed, *O. tsutsugamushi* seem to cluster at either spindle pole (Fig. 1C). This localisation suggests a potential interaction with the microtubule network to facilitate partitioning. In anaphase, the bacteria remain in proximity to the segregating chromosomes, although not tightly clustered at the spindle poles (Fig. 1C). Some individual bacteria are seen in the spindle midzone. By cytokinesis, *O. tsutsugamushi* are often visually equally distributed between daughter cells (Fig. 1C). To assess the generalisability of the results, we examined a second strain of *O. tsutsugamushi* (Karp), which displayed a similar mitotic localisation pattern (Supp. Fig. 1). This suggests that this behaviour may be conserved across strains, and that it potentially reflects a shared mechanism for intracellular positioning

Next, we evaluated bacterial inheritance during host cell division by quantifying the bacterial signal in each daughter cell during telophase or cytokinesis and calculated a symmetry index, where 0 indicates complete asymmetry and 1 represents perfect symmetry. Because a random distribution would tend toward greater symmetry with increasing bacterial load, total bacterial signal was included in the analysis. As a reference for typical organelle distribution, we assessed mitochondrial symmetry in uninfected RPE1 cells, which were synchronised for 16 hours, fixed, and labelled against the outer mitochondrial membrane protein TOM20. The mitochondrial signal in each daughter cell was quantified to calculate a symmetry index. Compared to mitochondria (mean symmetry index = 0.92), *O. tsutsugamushi* exhibited significantly less symmetrical distribution, yet generally showed a tendency toward symmetric partitioning (mean = 0.71; Estimate = –1.85, SE = 0.72, z = –2.52, p = 0.0096; Fig. 1D) and bacterial load was weakly positively correlated with the symmetry index (Estimate = 0.015, SE = 0.0057, z = 2.72, p = 0.0065). This relationship is also reflected in the scatter plot (Fig. 1E), which shows a slight but significant positive association between bacterial abundance and symmetry index (Spearman’s ρ = 0.39, p = 0.002), indicating that higher bacterial load is associated with slightly more equal partitioning. To understand when this symmetry is established, we next compared it across earlier mitotic stages. Compared to metaphase (mean = 0.81), the symmetry index did not differ significantly in anaphase (mean = 0.64; Estimate = −0.61, SE = 0.40, z = −1.53, p = 0.13), telophase (mean = 0.69; Estimate = −0.43, SE = 0.34, z = −1.26, p = 0.21), or cytokinesis (mean = 0.73; Estimate = −0.08, SE = 0.35, z = −0.22, p = 0.83), and bacterial load remained weakly positively associated with the symmetry index (Estimate = 0.0092, SE = 0.0041, z = 2.24, p = 0.025). This indicates that the near-symmetric distribution of *O. tsutsugamushi* is already established by metaphase, as soon as the bipolar spindle is formed. Although symmetry showed a weak positive correlation with bacterial load, this alone cannot account for the bias toward more balanced partitioning observed, especially given the non-random mitotic localisation of *O. tsutsugamushi.* Together, these findings support the presence of a regulated inheritance mechanism that promotes balanced partitioning.

The close association of *O. tsutsugamushi* with spindle poles, combined with previous evidence that the bacteria utilise microtubule-based transport suggests that their localisation and inheritance during host mitosis may rely on microtubules. Since *O. tsutsugamushi* cannot be genetically modified to disrupt microtubule interactions directly, we generated a stable RPE1 cell line expressing GFP-ScaC. This approach was designed to saturate ScaC-binding sites on BicD1/BicD2, thereby preventing the bacteria from engaging in microtubule-based transport. These cells were then synchronised and analysed 1 hour and 3 hours after release from mitotic arrest and compared to WT cells. In contrast to wild-type (WT) RPE1 cells (Supp. Fig. 2), GFP-ScaC overexpressing cells showed no strong spindle pole localisation following bipolar spindle formation (Fig. 2A). Additionally, *O. tsutsugamushi* visually appeared to be partitioned more asymmetrically (Fig. 2A) at cell division than in WT cells (Supp. Fig. 2). Compared to WT cells (mean symmetry index = 0.67), *O. tsutsugamushi* exhibited significantly less symmetrical distribution in GFP-ScaC overexpressing cells (mean = 0.44; Estimate = –0.90, SE = 0.29, z = - 3.12, p = 0.0018; Fig. 2B).

**Figure 2:**
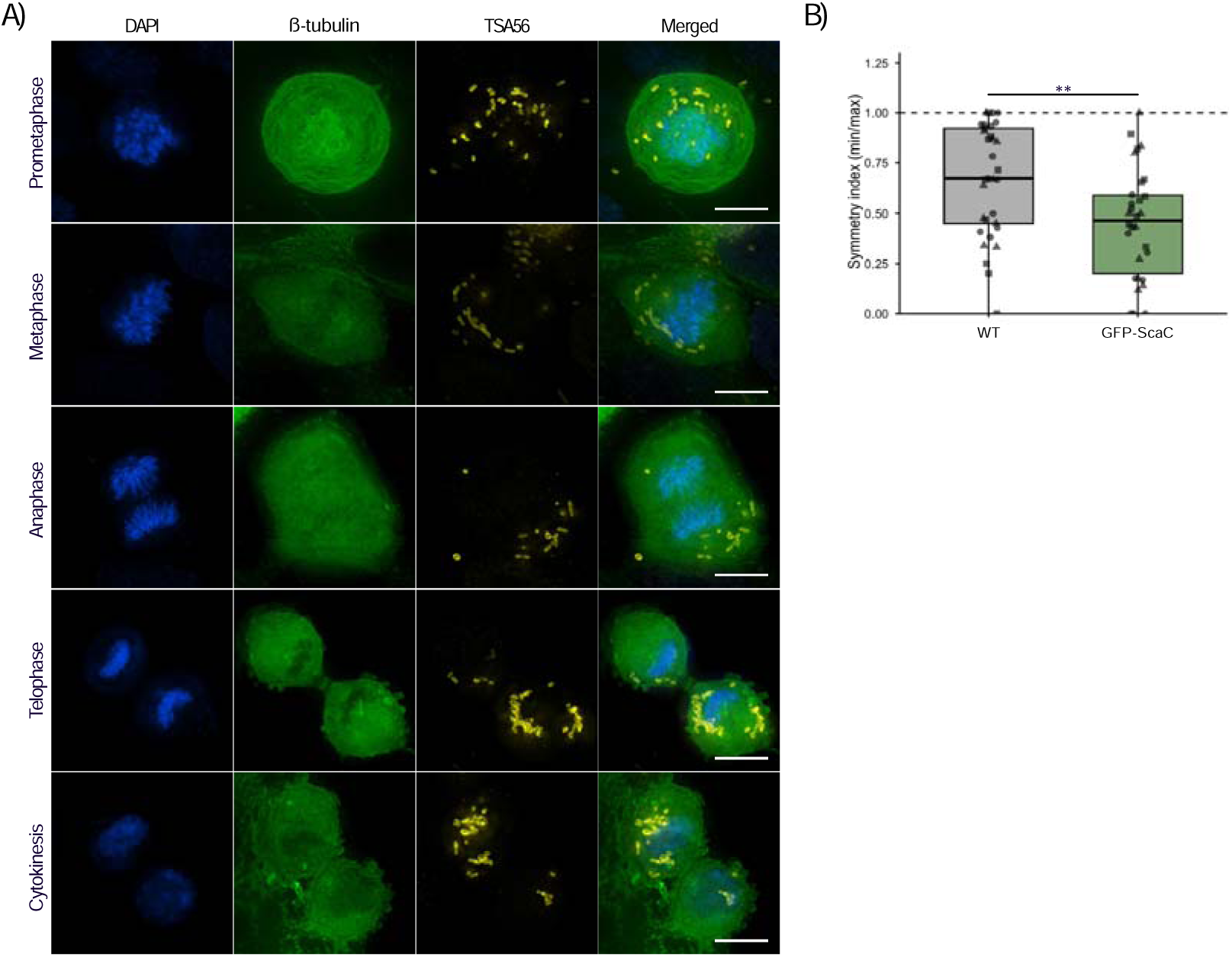
GFP-ScaC expression results in a reduced spindle pole localisation of *O. tsutsugamushi* in dividing host cells, as well as reduced symmetry in bacterial load per daughter cell. A. Representative image of GFP-ScaC Rpe1 cells infected for 6 days with Ot_TA686. Blue = DAPI (DNA), green = GFP-ScaC, yellow = anti-TSA56. Cells were synchronised using STLC, and samples were fixed 1 hour and 3 hours post-release. Images were acquired using a widefield microscope at 100X magnification. Scale bars represent 10 μm B. Quantification of symmetry index (min/max bacterial number in each daughter cell) of dividing cells in telophase or cytokinesis. A symmetry index of 1 indicates perfect symmetry. (GFP-ScaC: n=10,10,10 from 3 replicates; WT: n= 13, 10, 10 from 3 replicates). Boxplots show medians and interquartile ranges. Statistical analysis by beta regression comparing GFP-ScaC to WT, considering the total bacterial number per cell. Significance: ** p < 0.01.

Our findings suggest that dynein-mediated transport along microtubules may contribute to bacterial localization at spindle poles and to achieving more balanced partitioning. Interestingly, within the Rickettsiales, *Wolbachia* have been shown to associate closely with centrosomes during early *Drosophila* embryogenesis and to rely on microtubule-based trafficking for their symmetric distribution^16^. During interphase and prophase, *Wolbachia* move between sister centrosomes, likely via astral microtubules. This movement ceases during anaphase, and the bacteria remain tightly associated with separating centrosomes, resulting in symmetric segregation. Treatment with microtubule inhibitors disrupts both their movement and centrosomal association, demonstrating a clear reliance on intact microtubule structures^16^.

These observations highlight that at least two members of the Rickettsiales (*O. tsutsugamushi* and *Wolbachia*) rely on microtubules for symmetric segregation with *O. tsutsugamushi* forming direct contact with microtubules, in contrast to *Wolbachia*. In light of this, we next asked how another intracellular Rickettsiales, *R. rhipicephali*, which is not known to associate with microtubules, is localised during host cell division.

### *R. rhipicephali* strongly inhibits host mitosis and is randomly distributed in the few observed mitotic cells

In contrast to *O. tsutsugamushi*, infection with *Rickettsia rhipicephali* strongly reduced host mitosis. Only 2.9% of cells underwent mitosis versus 21.9% in the corresponding control (p = 1.73 × 10^-21^; Fig. 3A & 3B). Given this strong inhibition of cell division, ensuring partitioning into daughter cells may be less relevant for *R. rhipicephali*.

**Figure 3:**
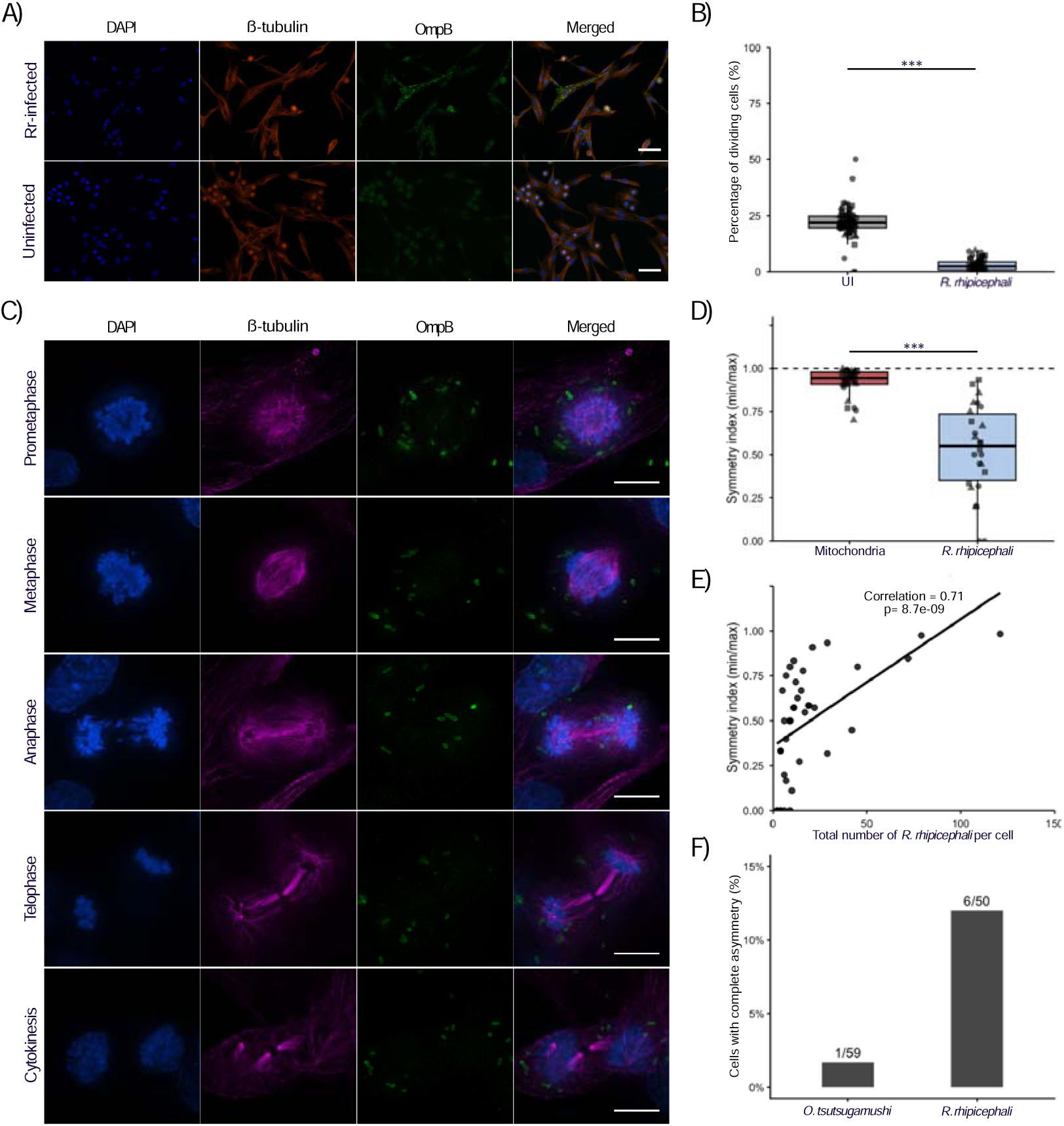
*R. rhipicephali* shows a random distribution pattern in mitotic host cells and are distributed asymmetrically between daughter cells A. Representative image of Rpe1 cells infected for 4 days with Rr. Blue = DAPI (DNA), red = anti-β-tubulin antibody, green = anti-OmpB. Images were acquired using a widefield microscope at 20X magnification. Scale bar = 50 μm. B. Quantification of data described in A. showing percentage of dividing cells in uninfected and infected samples. Boxplots display the median and interquartile range for each group. Each data point represents a single imaging field (n = 24 fields per replicate), acquired from three biological replicates. Given that an average of 93.46% ± 3.63% (SEM) of *Rickettsia*-infected were infected across the repeats, we can confidently attribute any observed changes in division frequency to the presence of the bacteria. C. Representative image of Rpe1 cells infected for 4 days with Rr. Blue = DAPI (DNA), pink = anti-β-tubulin antibody, green = anti-OmpB. Cells were synchronised with STLC and fixed 1 h and 3 h post-release. Images were acquired using a widefield microscope at 100X magnification. Scale bars = 10 μm. D. Quantification of symmetry index (min/max bacterial number or mitochondrial signal in each daughter cell) of dividing cells in telophase or cytokinesis. A symmetry index of 1 indicates perfect symmetry. (Mitochondria: n=12, 10, 9 from 3 replicates; Rr: n=10, 8, 8 from 3 replicates). E. Scatter plots showing the relationship between total bacterial load and symmetry index in dividing cells from metaphase and later mitotic stages. Each point represents a single cell. The black line indicates the fitted linear regression (Rr: n=27, 15, 11 from 3 replicates). F. Proportion of dividing Rpe1 cells exhibiting complete asymmetry in distribution of Ot_TA686 or Rr (symmetry index = 0) from metaphase and later mitotic stages. Complete asymmetry indicates that all bacteria were segregated into a single daughter cell. (Ot_TA686: n=12, 30, 3, 8, 6 from 5 replicates; Rr: n=27, 15, 11 from 3 replicates). Boxplots show medians and interquartile ranges. Statistical analysis by beta regression comparing *R. rhipicephali* to mitochondria, considering total bacterial number per cell. Significance: *** p < 0.001.

To visualise the intracellular distribution of *R. rhipicephali* during host mitosis, infected RPE1 cells were prepared as described with *Orientia*-infected cells. Bacteria were detected using antibodies against the outer membrane protein OmpB, and cells were also co-labelled with β-tubulin. In contrast to *O. tsutsugamushi*, *R. rhipicephali* displayed a random localisation pattern throughout mitosis without any clear enrichment at the spindle poles, midzone, or near chromosomes at any mitotic stage (Fig. 3C). This is consistent with the fact that *Rickettsia* species are not known to interact with microtubules and are therefore not expected to be associated with the mitotic spindle.

To assess how these localisation patterns affect bacterial inheritance, we measured the symmetry of distribution between daughter cells. Compared to mitochondria (mean symmetry index = 0.92), *R. rhipicephali* displayed an even greater degree of asymmetry than *O. tsutsugamushi* (mean = 0.53; Estimate = –2.83, SE = 0.68, z = –4.73, p = 2.2e-06; Fig. 3D) and bacterial load also positively correlated with the symmetry index (Estimate = 0.035, SE = 0.0086, z = 4.05, p = 5.2e-05). A direct correlation analysis revealed a stronger positive association between bacterial abundance and symmetry index for *R. rhipicephali* (Spearman’s ρ = 0.71, p = 8.7 × 10^-9^, Fig. 3E) than for *O. tsutsugamushi*. This suggests that, for *R. rhipicephali*, bacterial load may play a greater role in promoting more equal partitioning. Importantly, complete asymmetry (symmetry index = 0) was also more frequent for *R. rhipicephali*-infected cells, representing 12% of cells (6/50) after spindle pole formation compared to only 1.7% in *O. tsutsugamushi*-infected cells (1/59) (Fig. 3F). These observations are consistent with a more random partitioning of *R. rhipicephali* between daughter cells, as reflected by their visually irregular patterns, a high frequency of extreme asymmetry, and generally greater overall asymmetry in distribution between daughter cells.

In summary, this study shows that *R. rhipicephali* and *O. tsutsugamushi* exhibit different adaptations to the host cell cycle. While both bacteria appear to employ some degree of cell-cycle arrest, *R. rhipicephali* may rely primarily on this to ensure its maintenance within host cells and the establishment of a stable intracellular niche. This could reduce the selective pressure for tightly regulated partitioning mechanisms that would result in a more symmetric bacterial distribution between daughter cells. In contrast, a substantial fraction of *O. tsutsugamushi*– infected cells still divide, and therefore *O. tsutsugamushi* may benefit from mechanisms that ensure equal inheritance during host cell division. Like *Wolbachia pipientis*^15,16^, *O. tsutsugamushi* appears to interact with the microtubule network to achieve near-symmetric partitioning. This is consistent with the strong reliance of both bacteria on vertical transmission^18,19^. However, whereas *Wolbachia* resides within host-derived vacuoles^7^, *O. tsutsugamushi* is cytoplasmic^8^. Its intracellular lifestyle is more similar to that of mitochondria, making it a particularly interesting model to study bacterial endosymbioses. While this work was conducted in mammalian cells, future studies in arthropod hosts will be critical to assess whether the same mechanisms govern symbiont segregation in their natural cellular environment. Altogether, our findings represent an important step toward understanding how cytoplasmic endosymbionts adapt to host division and ensure long-term symbiotic stability.

## Materials and Methods

### Cell cultures

Human retinal pigment epithelial (RPE-1, ATCC, Cat# CRL-4000) and mouse fibroblast (L929, ATCC, Cat# CCL1) cell lines were cultured as monolayers in T25 or T75 flasks using Dulbecco’s Modified Eagle Medium (Gibco, Cat#41965062) supplemented with 10% heat-inactivated fetal bovine serum (Gibco, Cat# 16140071). Cells were maintained at 37 °C in a humidified incubator with 5% CO2. When cultures reached 90-100% confluence, cells were passaged using 0.25% Trypsin-EDTA (Gibco, Cat#25200056).

### Plasmid isolation

Escherichia coli DH5α containing the GFP-ScaC plasmid (GeneCopoeia; previously described^10^) was grown overnight in LB medium with 100 μg/mL ampicillin. Plasmid DNA was isolated using the QIAprep Spin Miniprep Kit (Qiagen) following the manufacturer’s instructions. Purified plasmid DNA was stored at –20°C until further use.

### Stable cell line generation via lentiviral transduction

To generate a stable RPE-1 cell line expressing GFP-ScaC, lentiviral particles were produced in Lenti-X 293T packaging cells (Takara Bio). Cells were co-transfected with three plasmids using Lipofectamine 3000 (Thermo Fisher Scientific): a transfer plasmid encoding GFP-ScaC (GeneCopoeia), a packaging plasmid (psPAX2, Addgene #12260), and an envelope plasmid (pMD2.G, Addgene #12259). The culture medium containing lentiviral particles was collected 48 hours post-transfection, filtered, and stored at −80 °C. RPE-1 cells were transduced with this viral supernatant for 48 hours to achieve stable integration. Lentiviral integration is random and results in permanent expression of the transgene.

### Bacterial stock preparation

*O. tsutsugamushi* strain TA686/Karp or *R. rhipicephali* were propagated in L929 mouse fibroblast cells cultured in 75-cm2 flasks. Uninfected L929 cells were maintained in Dulbecco’s Modified Eagle Medium (DMEM; Gibco, Cat# 41965-039) supplemented with 10% fetal bovine serum (FBS), at 37 °C in a 5% CO2 atmosphere. For infection, frozen stocks of bacteria were added to cells at 60–90% confluency. After infection, cultures were incubated in DMEM supplemented with 5% FBS at 35 °C. At six days post-infection, the culture medium was removed, and cells were washed once with fresh media. Infected cells were resuspended in fresh DMEM and detached using a cell scraper (Corning, Cat# CLS3010). The cell suspension was transferred to 1.5 mL microcentrifuge tubes and subjected to mechanical lysis using 0.1 mm glass beads (BioSpec, Cole-Parmer, Cat# 11079101) and a mini homogenizer (Fisherbrand, Cat# 15525799) operated at power at level 5 (150 × g; 5.0 m/s) for 1 minute. Lysates were centrifuged at 800 × g for 3 minutes to pellet glass beads and host cell debris. The resulting supernatant was transferred to new microcentrifuge tubes and centrifuged at 14,000 × g for 5 minutes to pellet the bacteria. The bacterial pellet was resuspended in SPG buffer (0.218 M sucrose, 3.76 mM KH2PO4, 7.1 mM K2HPO4, 4.9 mM monosodium L-glutamate). Bacterial concentrations were quantified by quantitative PCR (qPCR), and stocks were aliquoted and stored at −80 °C until use. For infection of RPE-1 cells, frozen bacterial stocks were thawed and used directly at a multiplicity of infection (MOI) of one host cell compared to 50 bacteria (1:50) for *O. tsutsugamushi* and a MOI of 1:75 for *R. rhipicephali*.

### Infection, synchronisation and drug treatment of RPE-1 cells

Experiments were set up in such a way as to fix the samples 6 days post infection for *O. tsutsugamushi* and 4 days post infection for *R. rhipicephali* to account for their differing growth rates and the earlier onset of host cell damage caused by Rickettsia replication ^4^.

Wild-type and GFP-ScaC-expressing RPE-1 cells were seeded into 25-cm2 flasks at a density of 2.8 × 105 cells per flask and incubated overnight at 37 °C in a 5% CO2 atmosphere. Cells were infected with a MOI of 1:50 for *O. tsutsugamushi* or 1:75 for *R. rhipicephali*. The culture medium was replaced 24 hours post-infection to remove unbound bacteria.

At 4 days post-infection for *O. tsutsugamushi* or 2 days post-infection for *R. rhipicephali*, cells were harvested and seeded into sterile μ-Slide 8-well ibiTreat chambers (Ibidi, Cat# 80806) at a density of approximately 0.3 × 10^5^ cells per well. Cells were incubated overnight to allow attachment. The following day, the medium was replaced with DMEM containing 5 μM S-trityl-L-cysteine (STLC, Alfa Aesar, Cat# L14384-06) to synchronise cells at mitosis, or an equivalent volume of dimethyl sulfoxide (DMSO, Sigma-Aldrich, Cat# D8418-100ML) for non-synchronised control samples. After 16–19 hours of incubation, the cells were washed three times with phosphate-buffered saline (PBS) and released into fresh DMEM for 1–3 hours to allow cell cycle progression before fixation for imaging.

Cells were washed with PBS and fixed in 4% paraformaldehyde (Thermo Fisher Scientific, Cat# 28906) for 15 minutes at 36°C. Fixed cells were permeabilised by sequential incubation in absolute ethanol (Thermo Fisher Scientific, Cat# 12478740) for 30 minutes on ice, followed by 0.5% Triton X-100 (Thermo Fisher Scientific, Cat# 85112) in PBS for an additional 30 minutes on ice. After permeabilisation, cells were washed three times with PBS.

For immunofluorescence microscopy of intracellular bacteria and host cell components, cells were incubated with primary antibodies overnight at 4°C in PBS supplemented with 1 mg/mL bovine serum albumin (BSA, Sigma-Aldrich, Cat# A9647-100G). Following primary antibody incubation, cells were washed three times with PBS and incubated with the corresponding secondary antibodies for 1 hour at 36°C in PBS containing 1 mg/mL BSA. Nuclei were counterstained with DAPI (1:1000 dilution). After labelling, the cells were maintained in mounting medium (20 mM Tris pH 8.0, 0.5% N-propyl-gallate, 90% glycerol).

Fig.1A, Fig.1C, Fig. 3A, Fig.3C, Supp. Fig. 1: For *O. tsutsugamushi* strain TA686, an in-house monoclonal human anti-TSA56 antibody was used (undiluted). For detection of *O. tsutsugamushi* strain Karp, a rat monoclonal antibody against TSA56 was used at a 1:200 dilution. *Rickettsia rhipicephali* was detected using an anti-OmpB antibody (Eurogentec, Cat# GB24035.F; 1:200). Host cell structures were labelled with a mouse monoclonal anti-β-tubulin antibody (Sigma-Aldrich, Cat# T4026; 1:500) for microtubules and a rabbit polyclonal anti-TOMM20 antibody (Sigma-Aldrich, Cat# P36970, 1:500) for mitochondria. Corresponding secondary antibodies used were anti-rat Alexa Fluor 488 (ThermoFisher, Cat# A-11006; 1:500), goat anti-human IgG Alexa Fluor 488 (Invitrogen, A-11013; 1:500), goat anti-rabbit Alexa Fluor 488 (ThermoFisher, Cat# A-11008; 1:500), and goat anti-mouse Alexa Fluor 555 (ThermoFisher, Cat# A28180; 1:500), and goat anti-rabbit IRDye 680RD (LICORbio, Cat# 926-68071).

Fig. 2A, Supp. Fig. 2: The bacterial antigen was detected using the in-house monoclonal human TSA56 antibody. For wild-type RPE-1 cells, the secondary antibody was goat anti-human IgG Alexa Fluor 488 and for GFP-ScaC-expressing cells, goat anti-human IgG Alexa Fluor 647 (Invitrogen, Cat# A-21445; 1:500) was used. Additionally, CellMask Deep Red (ThermoFisher, Cat# A-11006; 1:1000) was used to label the cell membrane.

### Immunofluorescence microscopy

For quantification of dividing cells and mitochondrial signal in mitotic cells, imaging was performed using a widefield fluorescence microscope (EVOS M7000, Invitrogen) with a 20X or a 40X objective lens controlled by the Invitrogen EVOS M7000 Imaging System. For quantification of dividing cells, the number of total nuclei was counted using CellProfiler (version 4.2.8). The nuclei were segmented using a global strategy (Minimum Cross-Entropy method). The number of mitotic cells were counted manually. The quantification of mitochondrial signal in mitotic cell was also performed using using CellProfiler. The host cell nucleus was segmented using a global strategy (Minimum Cross-Entropy method), in which the cell was defined as a secondary object correlated with the nucleus and segmented using an adaptive strategy (Otsu method) with an adaptive window of 50. The integrated intensity of the TOM20 signal (mitochondrial signal) was measured within the object cell.

All other imaging was performed using an Eclipse Ti2 widefield fluorescence microscope (Nikon) equipped with a 100x/1.4NA oil immersion objective lens and controlled by NIS-Elements AR 5.42.07 software. All images were acquired as Z-stacks to capture the full three-dimensional structure of the cells. Post-acquisition, images underwent 3D deconvolution to improve resolution and reduce out-of-focus light. Images were background-corrected by subtracting a constant intensity value from all channels and processed using maximum intensity projection (MaxIP). For the quantification of the symmetry index and the percentage of dividing bacteria, the bacteria were counted manually throughout all Z-stacks.

### Statistical methods

The fraction of dividing cells in infected versus uninfected (UI) samples was compared using two-sample Mann–Whitney tests. To compare the symmetry index of bacteria in dividing cells with that of mitochondria, a beta regression model in the style of an analysis of covariance was applied, using the total bacterial count or total mitochondrial signal as a covariate. The symmetry index was adjusted using a standard transformation for beta regression by applying the formula: symmetry index × (n – 1) + 0.5) / n, where n is the sample size. This ensured that all values fell strictly within the (0, 1) interval required by the beta regression model^20^. The same beta regression approach was used for comparing the symmetry index in the GFP-ScaC expressing cells to WT cells. Correlations between bacterial load and symmetry index were assessed using Spearman’s rank correlation. In all box plots, the middle line represents the median, and the lower and upper hinges indicate the first and third quartiles, respectively; whiskers extending out of the box indicate the variability beyond the upper and lower quartiles; data points beyond the whiskers are considered outliers. All statistical values reported on the figures are as follows: *p-value < 0.05; **p-value < 0.01; ***p-value < 0.001. All data were analysed using R version 2023.12.1+402.

## Acknowledgements

We would like to thank members of the Salje lab from the Cambridge Institute for Medical Research, University of Cambridge, UK: Rachel Jackson for providing the cell line; Beth Harris and Magda Rogowska-van der Molen for bacterial stocks; Panjaporn Chaichana for the antibody against TSA56; and Fran Aylward and Owen Somhorst for reagents. This work was supported by a Wellcome Trust Senior Research Fellowship to JS (224277/Z/21/Z).

## Notes

### Competing Interest Statement

The authors have declared no competing interest.

