## Supplementary Material for "Symmetric Inheritance of the Cytoplasmic Bacterium *Orientia tsutsugamushi* During Mitosis"

**Supplementary Figure 1: *O. tsutsugamushi* strain Karp displays similar localisation to *O. tsutsugamushi* strain TA686.** Representative images of Rpe1 cells infected for 6 days with Ot_Karp. Blue = DAPI (DNA), pink = anti-β-tubulin antibody, green = anti-TSA56. Cells were synchronised with STLC and fixed 1 h and 3 h post-release. Images were acquired using a widefield microscope at 100X magnification. Scale bars = 10 μm.

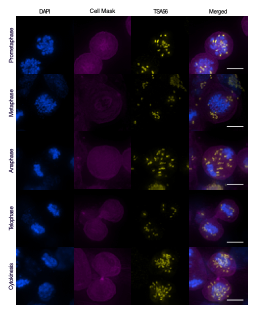
**Supplementary Figure 2: O. tsutsugamushi in dividing wild-type (WT) RPE-1 grown/infected/synchronised in parallel to GFP-ScaC cells show the same spindle pole localisation as described in Fig. 3.** Representative images of WT Rpe1 cells infected for 6 days with Ot_TA686. Blue = DAPI (DNA), pink = Cell mask, yellow = anti-TSA56. Cells were synchronised with STLC and fixed 1 h and 3 h post-release. Images were acquired using a widefield microscope at 100X magnification. Scale bars = 10 μm.
